# Local trait responses and aquatic microclimates increase projected habitat suitability for field-collected *Anopheles stephensi* in southern Ethiopia

**DOI:** 10.64898/2026.07.21.739549

**Authors:** Paul J. Huxley, Maxwell G. Machani, Dawit Hawaria, Samuel S.C. Rund, Guiyun Yan

## Abstract

**Background:** Trait-based models of mosquito environmental suitability commonly use global thermal performance curves compiled from laboratory studies. These curves may misrepresent suitability when local populations or larval habitats differ from globally synthesised expectations. This issue is particularly relevant for *Anopheles stephensi*, an urban malaria vector expanding in the Horn of Africa, where artificial aquatic habitats may generate microclimates poorly represented by global trait datasets or gridded air-temperature products.

**Methodology/Principal Findings:** We compared juvenile survival, development, and projected maximal population growth rate, *r*_m_, using locally measured data from field-collected Ethiopian *An. stephensi* and a global comparator combining published juvenile trait data. We fitted thermal performance curves for juvenile survival and development, used these curves to estimate temperature-dependent *r*_m_ as a metric of habitat suitability, and projected *r*_m_ across two matched temperature inputs: logger-measured larval habitat water temperature and ERA5-Land 2 m air temperature. Local juvenile trait responses produced higher projected *r*_m_ than the global comparator across monitored habitats. Measured aquatic temperatures were warmer than ERA5-Land air temperatures, with a mean daily water–air offset of 3.26*^◦^*C, and were more spatially heterogeneous among habitats. ERA5-Land resolved only two unique temperature series across the six monitored habitats. Sensitivity analyses showed that the Local–Global difference was robust for logger-measured water temperatures, whereas the ERA5-based difference was partly amplified by cooler temperatures below the local experimental range.

**Conclusions/Significance:** This study provides the first locally derived juvenile thermal performance curves and temperature-dependent population growth estimates for field-collected *An. stephensi* in Africa, demonstrating that locally measured juvenile traits consistently predict higher habitat suitability than globally synthesised trait data. These findings highlight the importance of incorporating local mosquito trait data and aquatic microclimate measurements into predictive models to improve assessments of *An. stephensi* establishment, spread, and malaria risk in newly invaded urban environments.

**Author Summary:** *Anopheles stephensi* is an invasive malaria mosquito that is rapidly spreading in urban areas of Africa. Predicting where this species can establish is difficult because many models use mosquito temperature-response data from long-maintained laboratory colonies rather than from recently collected African populations. We measured juvenile survival and development in *An. stephensi* collected from Ethiopia, and compared projections based on these local data with projections based on published laboratory-colony data. We also compared field-measured larval habitat water temperature with ERA5-Land air temperature. We found that local juvenile trait responses produced higher projected population growth than the global comparator across monitored habitats. Field-measured water temperatures were warmer and more variable among habitats than gridded air temperatures, which reduced habitat-level differences. These findings show that local mosquito biology and larval habitat microclimate measurements can change fine-scale estimates of suitability. This matters for surveillance and control because urban water containers may create suitable conditions that are poorly represented by broad climate datasets.

## Introduction

Environmental suitability for mosquito populations is shaped by the combined effects of climate, habitat availability, and mosquito life-history traits. Mosquitoes are ectotherms, so temperature has a particularly strong influence on development, survival, fecundity, biting, and pathogen transmission. Trait-based thermal performance curves (TPCs) are widely used to translate temperature into estimates of vector population growth or transmission potential [16, 24, 37]. These approaches have been valuable because they provide mechanistic links between climate and vector biology, rather than relying only on correlations between observed distributions and environmental covariates.

A common implementation of this framework is to use globally synthesised trait curves [35, 40]. In practice, such curves are often built by pooling data across studies, populations, laboratory colonies, experimental conditions, and geographic origins. This aggregation is often necessary because, despite the growing use of mechanistic models for estimating vector suitability, *R*_0_, transmission potential, and geographic distribution, temperature-dependent life-history trait data remain sparse for many vector species, including *Anopheles stephensi* mosquitoes. Pooling therefore increases taxonomic and thermal coverage and allows models to be parameterised where local data are unavailable. However, it also assumes that the resulting average trait response is an adequate proxy for the populations and environments being projected. That assumption may be reasonable for broad comparative analyses, but it is less secure when the objective is to infer suitability for a specific invasion front, field site, or local habitat type. Local populations may differ from global expectations because of genotype [5], colony history [15], larval habitat conditions [16–18], acclimation, experimental protocol, or adaptation to local climatic regimes [9, 10].

This issue is especially relevant for *An. stephensi*. Although the species is historically associated with South Asia and parts of the Middle East, it has recently expanded in Africa and is now a major concern for urban malaria transmission in the Horn of Africa [38, 43]. Recent genomic work suggests that the African invasion originated from South Asian source populations, with Djibouti acting as a bridgehead population that subsequently seeded multiple invasion fronts across East Africa and Yemen [11]. This invasion history is important because most available trait-based studies of *An. stephensi* rely on long-established laboratory colonies originally derived from South Asian or Middle Eastern populations. Care is therefore needed when extrapolating these responses to recently invaded African populations: invasive populations may have passed through founder events, bottlenecks, or subsequent ecological filtering, while laboratory colonies can diverge from wild populations through colonisation effects, laboratory adaptation, and genetic drift [2, 13, 22, 34]. At the same time, *An. stephensi* has been reported from African settings with temperatures substantially warmer than those typical of parts of its native range [33, 42]. The species also breeds readily in artificial water-storage containers, allowing persistence in urban and peri-urban environments where aquatic habitat temperatures can depart from standard gridded air-temperature products [7, 25, 28]. Taken together, invasion history, laboratory-colony provenance, and aquatic habitat microclimates all suggest that projections based only on globally averaged thermal performance curves or gridded air-temperature data may misrepresent fine-scale habitat suitability in the invasive range of *An. stephensi*.

Juvenile traits are central to this problem because they determine recruitment into the adult population. Juvenile survival, *p*_EA_, and development time, *α*, jointly regulate the rate at which individuals become adults and therefore strongly influence projected maximal population growth rate, *r*_m_ [17, 29]. If local juveniles survive better or develop faster at the temperatures experienced in field habitats than predicted by global TPCs, then global curves will underestimate population suitability. Conversely, if global curves overstate juvenile performance, they will overestimate suitability. Either direction matters for interpreting invasion and transmission risk, seasonal population dynamics, and the potential productivity of local aquatic habitats.

Here, we test whether local juvenile trait responses alter projections of *An. stephensi* habitat suitability in southern Ethiopia, and whether the temperature source used for projection changes inferred suitability. We compare two juvenile trait parameterisations: a local dataset measured from field-collected Ethiopian mosquitoes and a global comparator combining published *An. stephensi* juvenile trait data. We fit survival and development-time TPCs to each dataset and evaluate projected *r*_m_ across two matched environmental inputs: field-measured larval habitat water temperatures and ERA5-Land 2 m air temperature extracted for the same sites and period. We ask whether globally derived trait responses systematically differ from local trait responses, whether those differences lead to under- or overestimation of habitat suitability, whether gridded air-temperature products preserve or obscure fine-scale variation in larval habitat thermal conditions, and whether the inferred Local–Global difference is robust to the limited thermal range of the local juvenile experiments.

## Materials and methods

We compared local and global juvenile trait parameterisations for *An. stephensi*. For each parameterisation, we fitted Bayesian thermal response models to juvenile survival and juvenile development time. We then used posterior predictions from these fitted trait models to estimate temperature-dependent maximal population growth rate, *r*_m_, across both a smooth temperature grid and observed field habitat temperature time series. We also extracted ERA5-Land 2 m air temperature data at the field sites to evaluate how projections changed when using a gridded air-temperature product rather than measured aquatic habitat temperature. The goal was not to build a complete distribution model for *An. stephensi*, but to isolate how differences in juvenile trait thermal responses translate into projected habitat suitability.

### Experimental setup

*Anopheles* larvae (F0) were collected from urban breeding habitats, including swimming pools, brick-making pits*/*pools, and water storage containers (Fig. 1) in Hawassa, southern Ethiopia (Fig. 1). Emerged adults were maintained under laboratory conditions and blood-fed to generate F1 progeny. Ovipositing females were identified morphologically as *An. stephensi* using standard taxonomic keys [8], and a subset was confirmed by PCR amplification of the internal transcribed spacer 2 (ITS2) region [3]. To characterise thermal conditions experienced by juveniles in the field, water temperatures were recorded in productive *An. stephensi* larval habitats during July 2025 using HOBO MX1102 data loggers (Onset Computer Corporation, Bourne, MA, USA). Data loggers were placed directly within breeding habitats and programmed to continuously record water temperature over a two-week period (Fig. 1).

**Figure 1:**
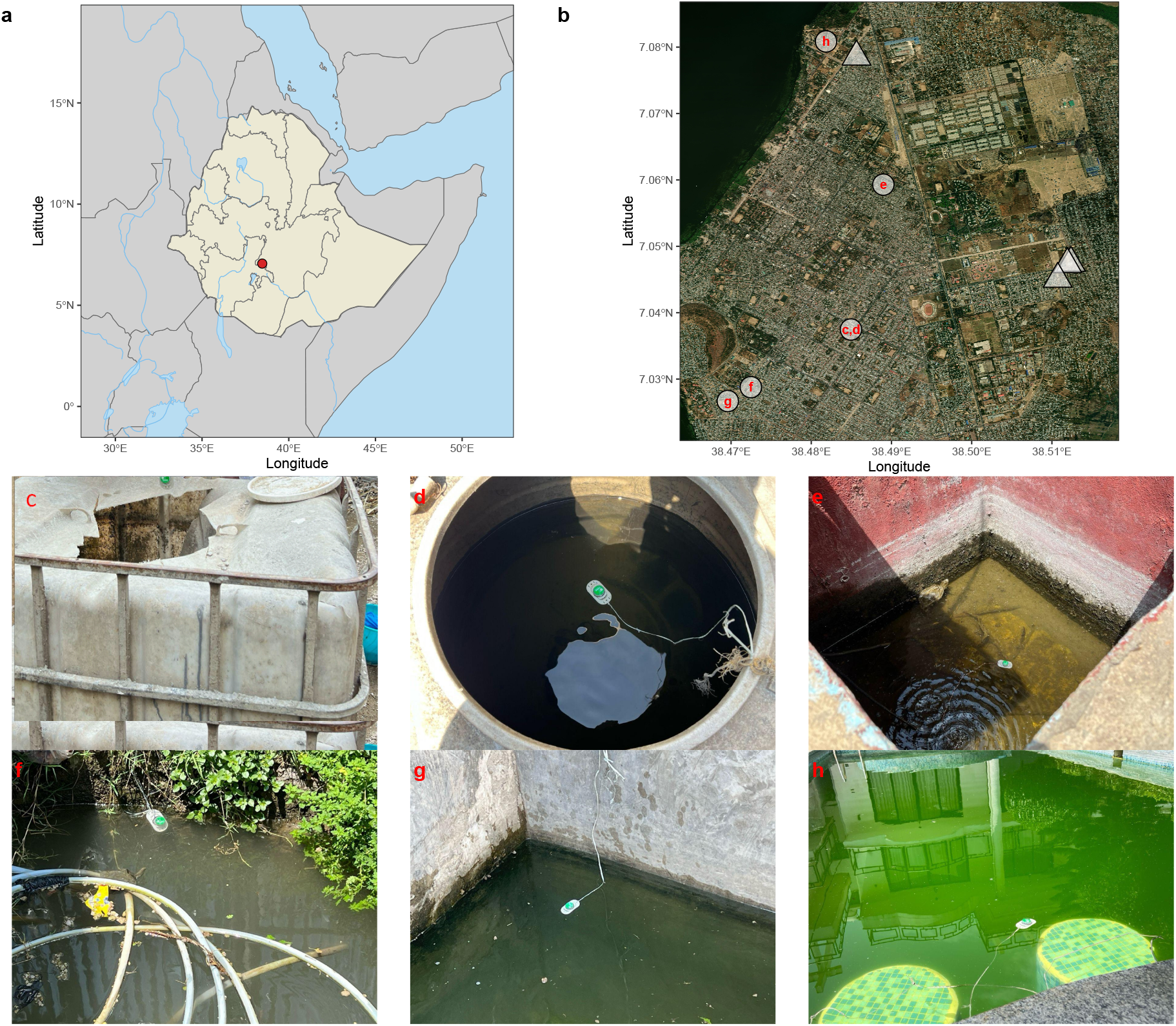
Field collection sites and temperature-monitored larval habitats in Hawassa, southern Ethiopia. (a) Country-level context map showing Ethiopia and the location of Hawassa. (b) High-resolution satellite/aerial basemap of the Hawassa sampling area showing georeferenced larval habitat collection sites. Lettered points correspond to the temperature-monitored habitats shown in panels c–h and were used in the population-growth projections; triangles indicate additional collection sites not used in the projection analysis. (c, d) Water storage tanks. (e) Water storage pool in a carwash area. (f, g) Water-holding pools at brick-making sites. (h) Swimming pool in a residential area. Site photographs: MGM; Basemap imagery: Esri World Imagery.

To estimate juvenile survival and development time, first-instar F1 larvae (<24 h post-hatching) were exposed to four constant temperature treatments (19°C, 26°C, 30°C, and 34°C). These temperatures were selected to represent the lower and upper limits of field-observed thermal conditions as well as projected warming scenarios. The 19°C treatment represented cooler seasonal temperatures, whereas 34°C simulated potential thermal stress associated with future climate warming. For each temperature treatment, 50 larvae were placed in a plastic container containing 1,000 mL of tap water. Each temperature treatment was replicated six times using freshly hatched larvae for each replicate. Water temperature was maintained using aquarium heaters equipped with thermostatic controllers. Water volume was maintained at 1,000 mL throughout the experiment to ensure consistent larval density and habitat conditions across treatments. Larvae were fed daily with equal amounts of finely ground commercial fish food (Tetramin®) to standardise resource availability across temperature treatments. All experimental units were maintained under standardised environmental conditions of 75% relative humidity and a 12:12 h light-dark photoperiod with dawn and dusk transitions. Larvae were monitored daily, and development time from hatching to adult emergence was recorded for each individual.

### Juvenile trait datasets

The local dataset consists of juvenile survival and development observations from the laboratory experiments conducted on *An. stephensi* collected from the Hawassa field sites shown in Fig. 1. These mosquitoes therefore represent a recently field-collected Ethiopian population rather than a long-maintained laboratory colony. The analysis retained local temperatures of 19, 26, 30, and 34*^◦^*C. Survival was summarised by replicate as the number of individuals surviving to emergence out of the initial cohort size. Development time was summarised by replicate as the emergence-weighted mean development time, calculated from daily male and female emergence counts.

The global comparator combined three externally derived sources that are all based on lab-maintained colonies. First, we used published *An. stephensi* trait data from Paaijmans et al. [27], including juvenile-to-adult survival and mosquito development rate records. These experiments used *An. stephensi* Liston mosquitoes cultured at Pennsylvania State University since 2008. The experiments were conducted at 90% relative humidity. Development rates were converted to development time as *α* = 1*/*development rate. Second, we included juvenile survival and development datasets from Huxley et al. [16], using the 75% relative humidity treatment to match the relative humidity used in the local Hawassa juvenile experiments. This study used an *An. stephensi* urban-type strain acquired from a longstanding (∼40 year) colony at the Walter Reed Army Institute of Research via the University of Georgia. Third, we included development and survival data from the high-food treatments in Tuno et al. [39]. These data used *An. stephensi* SDA-500, a strain obtained from the London School of Hygiene & Tropical Medicine and originally derived from Pakistan, then maintained for several hundred generations in Japan. The relative humidity associated with the Tuno et al. data was measured intermittently across a range of 60-80% (Personal communication).

Relative humidity was recorded because recent work has shown that humidity can modify thermal responses in *An. stephensi* [4, 16, 19]. The global comparator is therefore not humidity-standardised: it includes externally derived trait observations measured under known 90% and 75% relative humidity conditions, plus one source for which relative humidity ranged between 60%-80%. The global comparator also differs from the local dataset in colony history and geographic provenance, combining long-maintained laboratory strains rather than recently field-collected African mosquitoes. The trait data from all three sources were folded into the global comparator rather than analysed as separate strain- or humidity-specific scenarios, so that the final analysis compared only *Local* versus *Global* juvenile trait responses. This global comparator is therefore best interpreted as an externally derived, heterogeneous laboratory-colony response rather than a locally fitted Hawassa response.

### Thermal performance models

For each scenario, we fitted juvenile survival and development-time models separately. Juvenile survival was modelled as a binomial response, with the number surviving to emergence as the response and the initial number of larvae as the binomial denominator. We fitted two candidate survival models: a quadratic binomial generalized linear model and a LINEX-style binomial thermal response model [21]. For each scenario, the survival model used for prediction was selected by Watanabe–Akaike information criterion (WAIC). If one model failed to return a valid WAIC, the other model was used; if both were unavailable or tied, the quadratic model was used by default.

Development time was modelled as a temperature-dependent continuous response using an exponential function:

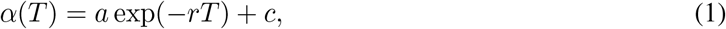

where *α*(*T* ) is juvenile development time at temperature *T* , and *a*, *r*, and *c* are fitted parameters. Models were fitted in R using `bayesTPC` and `nimble` [32, 36]. Posterior predictions were generated across the temperature range represented in the field habitat data.

### Population growth projection

We projected maximal population growth rate, *r*_m_, using an analytic approximation based on juvenile development time and juvenile survival. Following a previously published demographic framework [6, 16], and using emergence probability directly, *r*_m_ was calculated as:

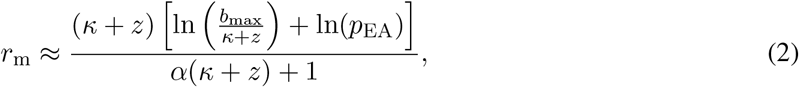

where *α* is juvenile development time, *p*_EA_ is juvenile-to-adult survival probability, *z* is adult mortality, *b*_max_ is maximum fecundity rate, and *κ* is the fecundity loss schedule. Parameter definitions are provided in Table 1. Adult mortality and fecundity were held constant across local and global scenarios so that differences in projected *r*_m_ reflected juvenile trait differences only. We used *z* = 0.12 day*^−^*^1^, *b*_max_ = 11.37 day*^−^*^1^, and *κ* = 0.01 day*^−^*^1^.

**Table 1:** Definitions of model parameters.

| Parameter | Units | Description |
| --- | --- | --- |
| $r_m$ | $\text{day}^{-1}$ | Maximal population growth rate |
| $\alpha$ | days | Juvenile-to-adult development time |
| $b_{\max}$ | $\text{eggs} \times \text{female}^{-1} \text{ day}^{-1}$ | Maximum fecundity rate |
| $\kappa$ | $\text{day}^{-1}$ | Fecundity loss rate |
| $z$ | $\text{day}^{-1}$ | Adult mortality rate |
| $p_{\text{EA}}$ | probability | Survival averaged across juvenile stages |

Posterior uncertainty was propagated by evaluating survival and development-time curves for each posterior draw at each temperature, calculating *r*_m_ for that draw, and then summarising the resulting *r*_m_ distribution. Because survival and development were fitted as separate models, this propagation assumes independent posterior uncertainty between traits. Habitat temperature observations were treated as fixed observed inputs; we did not propagate measurement error in logger temperatures, uncertainty in ERA5-Land estimates, or additional stochastic variation in habitat temperature beyond the recorded time series.

### Field sites and habitat temperature monitoring

Field collections and habitat temperature monitoring were conducted in Hawassa, Southern Ethiopia. Larval habitat collection sites were georeferenced in the field, giving 12 mapped aquatic habitat locations. Of these, six habitats were instrumented with temperature loggers and were used for the habitat suitability projections. The remaining georeferenced habitats are shown as additional collection sites but were not included in the temperature time-series projections.

We used measured aquatic habitat temperatures as the primary microclimate input because juvenile mosquito traits are expressed in aquatic habitats rather than in free-air conditions. This distinction is important because aquatic larval habitat temperatures can differ substantially from adjacent air temperatures, with consequences for estimated immature development and population growth [28]. To evaluate how a gridded air-temperature input would alter projected suitability, we also extracted ERA5-Land 2 m air temperature from the Copernicus Climate Data Store for the same monitored habitats and observation period. ERA5-Land values were converted from Kelvin to degrees Celsius and matched to the logger data by site and hour.

Field sites were mapped in R using `sf` [30, 31], `ggplot2` [41], `ggspatial` [12], and `rnaturalearth` [23]. The country-level panel shows Ethiopia with major physical features and the location of Hawassa, while the local panel shows the georeferenced larval habitat collection sites. Sites included in the projection analyses are distinguished from the remaining collection sites using different letters and symbols in Fig. 1.

### Field habitat temperature projections

We evaluated local and global trait projections across two matched temperature inputs: measured aquatic habitat temperatures from the six temperature-monitored larval habitats shown in Fig. 1, and ERA5-Land 2 m air temperature extracted for the same habitat coordinates and period. This produced projections for each combination of trait parameterisation and temperature source: *Local* traits with logger habitat water temperature, *Global* traits with logger habitat water temperature, *Local* traits with ERA5-Land 2 m air temperature, and *Global* traits with ERA5-Land 2 m air temperature.

For each temperature observation, we calculated posterior draws of *r*_m_ under both trait parameterisations. We then summarised suitability at two levels. First, for each habitat, date, and temperature source, we averaged *r*_m_ across within-day observations separately for each posterior draw and summarised the resulting draw-level daily means. Second, for each habitat and temperature source, we calculated mean daily *r*_m_ across the observation period for each posterior draw, then summarised posterior medians and 95% credible intervals. Because multiple habitat coordinates fell within the same ERA5-Land grid cell, extracted ERA5-Land temperature series were not always unique among habitats.

### Sensitivity analyses

Local juvenile trait data were available only from 19–34*^◦^*C, so we performed sensitivity analyses to assess whether projected Local–Global differences were driven by extrapolation beyond the local experimental range or by the broader thermal support of the global comparator. First, we quantified the proportion of logger-measured water temperatures and ERA5-Land 2 m air temperatures that fell within the local experimental range. We then repeated the habitat suitability summaries after restricting projection temperatures to observations within 19–34*^◦^*C. This tested whether Local–Global differences persisted when temperatures outside the local experimental support were excluded.

Second, we refitted the global comparator using only global juvenile trait observations within the local experimental temperature window. This restricted global fit was projected onto the same temperature inputs and compared against the original Local and Global projections. This analysis tested whether the broader thermal support of the global dataset, and the resulting more strongly unimodal fitted curve shape, contributed to lower projected suitability under the original global comparator. These sensitivity analyses were treated as diagnostic checks rather than as replacements for the main analysis, because excluding thermal observations from the global comparator removes information about its lower and upper thermal responses.

Full sensitivity-analysis summaries are reported in Supplementary Tables S1-S3 and Supplementary Figs. S1-S2.

## Results

### Local and global juvenile trait responses differ across temperature

Observed juvenile survival and development-time data differed clearly between the local and global trait datasets (Fig. 2). The local dataset was concentrated across the field-relevant temperature range of 19–34*^◦^*C, whereas the global comparator combined a wider and more heterogeneous set of externally derived observations, including literature-derived juvenile trait data and the 75% RH juvenile dataset. These differences were reflected in the fitted thermal performance curves (TPCs).

**Figure 2:**
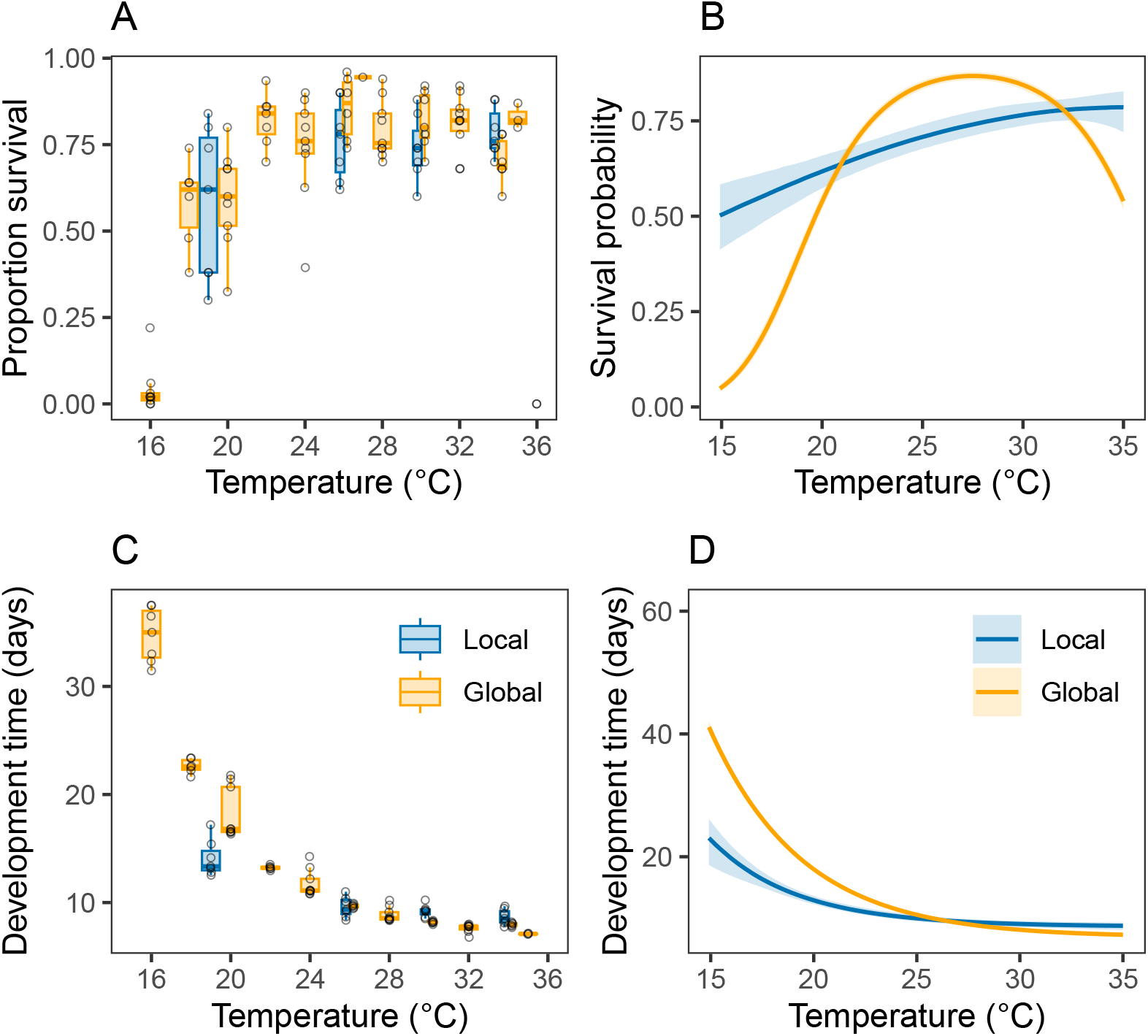
Observed and fitted juvenile trait responses for local and global *Anopheles stephensi* datasets. Observed juvenile survival and development-time summaries are shown alongside posterior-predicted thermal performance curves. Panels show observed survival (A), predicted survival TPCs (B), observed development time (C), and predicted development-time TPCs (D). Points and boxplots show replicate-level observations; lines show posterior medians; ribbons show 95% credible intervals.

For juvenile survival, the global comparator produced a strongly unimodal response, with low predicted survival at cooler temperatures, high survival at intermediate temperatures, and declining survival toward the warm end of the fitted range. In contrast, the local survival curve was flatter across the observed temperature range and did not show the same pronounced high-temperature decline. Development-time responses also differed between datasets. The global comparator predicted substantially longer development times at cooler temperatures and a steep decline in development time with increasing temperature, whereas the local curve predicted shorter development times across much of the field-relevant temperature range and a less extreme cool-temperature penalty. Together, these fitted trait responses indicate that local and global juvenile datasets translate the same environmental temperatures into different estimates of juvenile performance.

### Measured aquatic microclimates were warmer and more spatially heterogeneous than ERA5-Land air temperature

Measured larval habitat water temperatures were consistently warmer than matched ERA5-Land 2 m air temperature (Fig. 3). Across 102 matched habitat-days, the mean water–air offset was 3.26*^◦^*C (median 3.13*^◦^*C; range 0.96–6.34*^◦^*C). Habitat-specific mean offsets ranged from 2.0*^◦^*C at the carwash pool to 5.0*^◦^*C at the second brick-making pool. Thus, the gridded 2 m air-temperature input systematically underestimated the thermal conditions measured directly within larval aquatic habitats.

**Figure 3:**
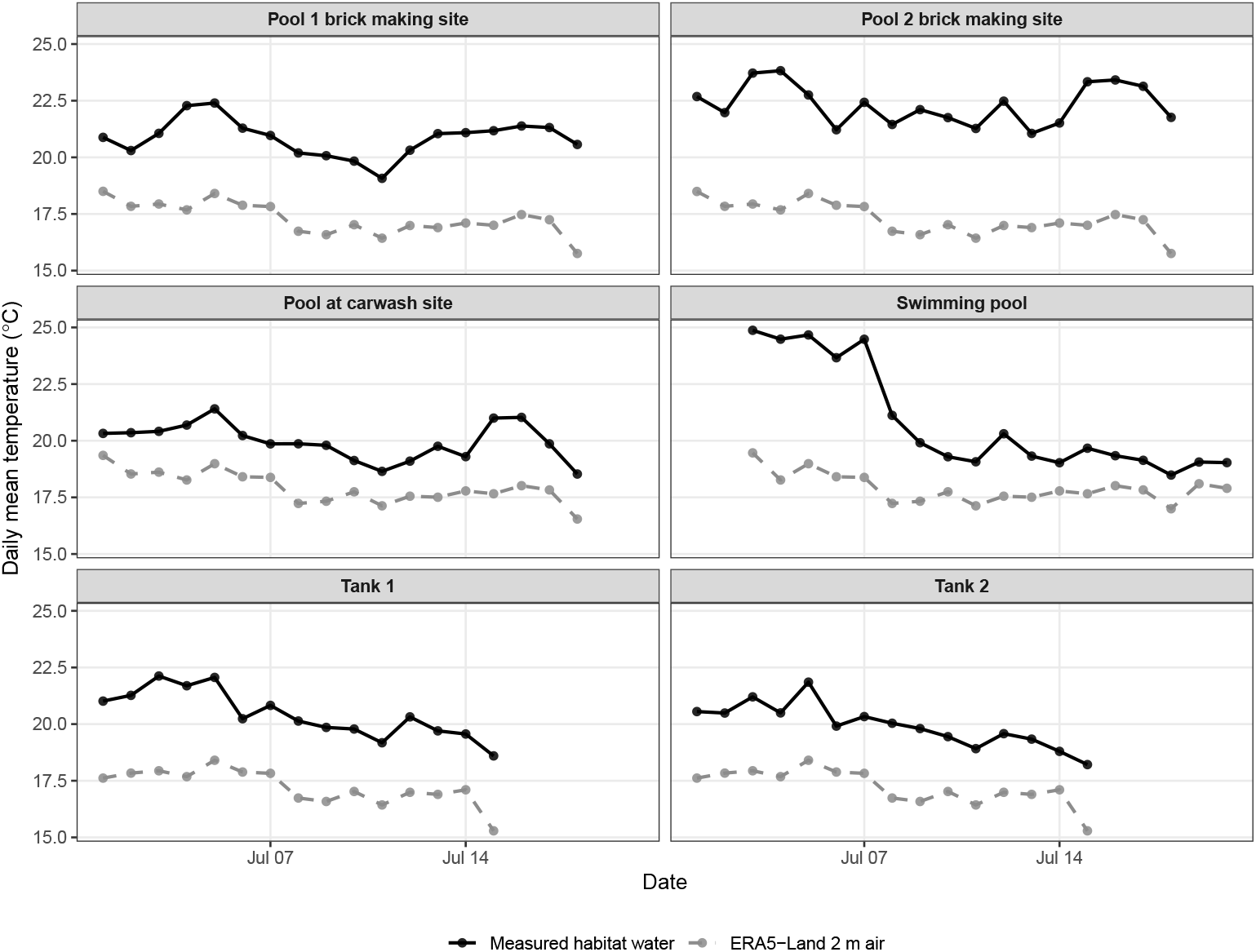
Daily mean measured habitat water temperature and ERA5-Land 2 m air temperature across monitored larval habitats. Lines show daily mean temperature for field-measured aquatic habitat water and matched ERA5-Land 2 m air temperature. Facets show the six temperature-monitored larval habitats used in the projection analysis.

The two temperature inputs also differed in their spatial resolution of habitat-level thermal variation. Logger measurements produced distinct aquatic temperature time series among habitats, whereas ERA5-Land resolved only two unique hourly temperature series across the six monitored habitats. Sites 1, 2, 4, and 5 shared one identical ERA5-Land series, while sites 3 and 6 shared a second identical series. Consequently, ERA5-based suitability projections were identical or near-identical for several habitats, whereas logger-based projections retained habitat-specific thermal differences. This indicates that gridded air-temperature products can smooth over fine-scale aquatic habitat heterogeneity relevant to immature mosquito develop-ment and survival.

### Global juvenile trait curves underestimate projected population growth across field habitats

Differences in juvenile trait responses translated into clear differences in projected maximal population growth rate, *r*_m_ (Fig. 4). Across the temperature grid, the local and global parameterisations produced dis-tinct *r*_m_ thermal performance curves. The global comparator produced a more peaked curve with higher pro-jected *r*_m_ near its warm-temperature optimum, but projected substantially lower *r*_m_ at cooler-to-intermediate temperatures. In contrast, the local parameterisation produced higher projected *r*_m_ across much of the temperature range most relevant to the sampled Hawassa aquatic habitats.

**Figure 4:**
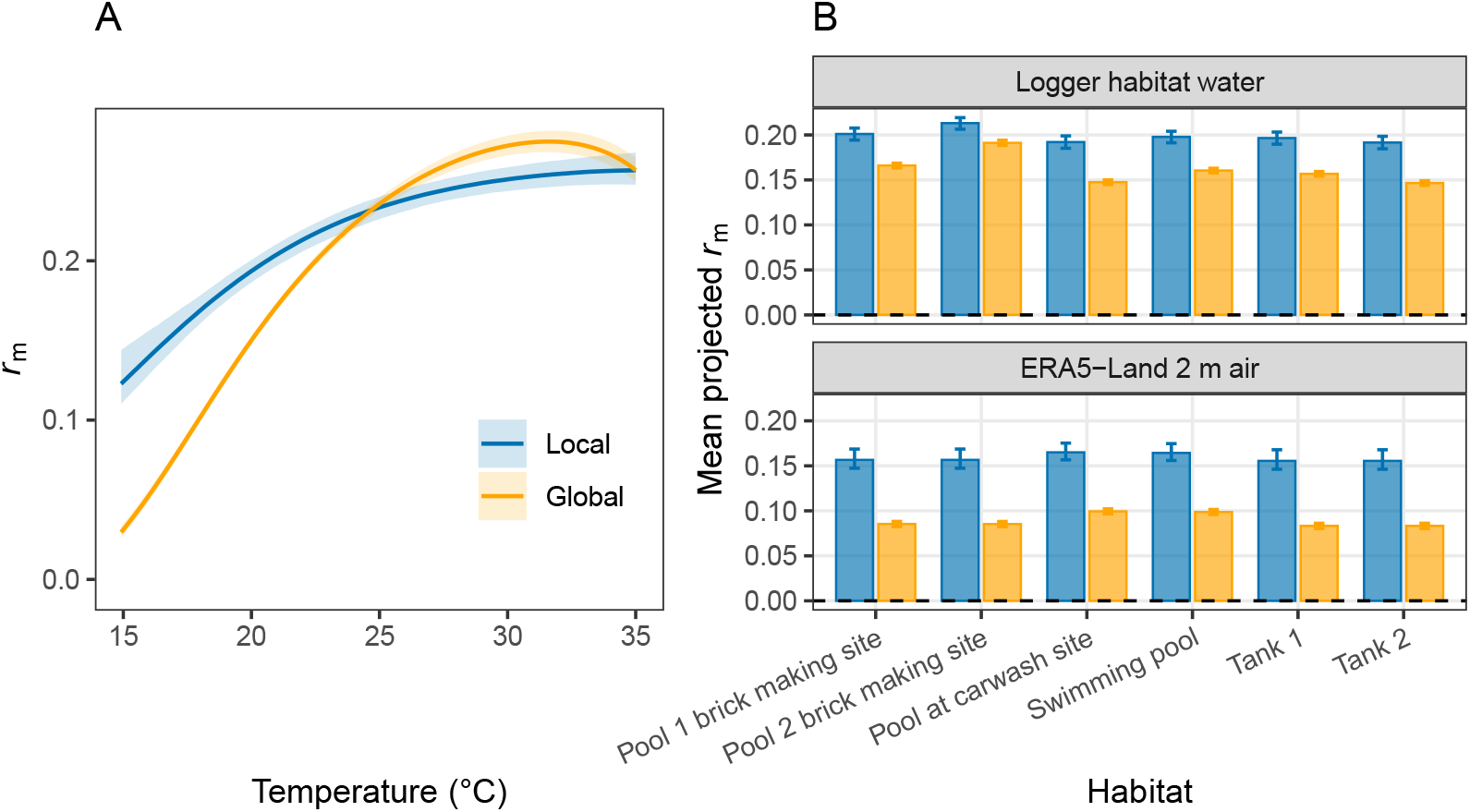
Local juvenile trait responses increase projected maximal population growth rate relative to global curves. (A) Temperature-dependent projections of maximal population growth rate, *r*_m_, generated by propagating juvenile survival and development-time posterior predictions through Eqn. 2. Lines show posterior medians and ribbons show 95% credible intervals. (B) Mean projected *r*_m_ by field habitat. Bars show posterior medians and error bars show 95% credible intervals.

When projected onto observed field habitat temperatures, this difference produced a consistent pattern of underestimation by the global comparator. Mean projected *r*_m_ was higher under the local trait parameter-isation than under the global parameterisation in all six sampled habitats (Fig. 4B). The magnitude of the difference varied among habitats, reflecting differences in habitat temperature distributions, but the direc-tion of the effect was consistent. Thus, although the global *r*_m_ TPC reached high values near its optimum, it underestimated habitat suitability across the temperatures actually experienced in the field.

### A Differences in projected suitability persist through time

Daily suitability projections showed that the divergence between local and global parameterisations per-sisted across the field temperature time series (Fig. 5). In every habitat, the local parameterisation produced higher daily mean *r*_m_ than the global comparator throughout the observation period. The absolute values and temporal fluctuations differed among habitats, but the local projection remained consistently above the global projection.

**Figure 5:**
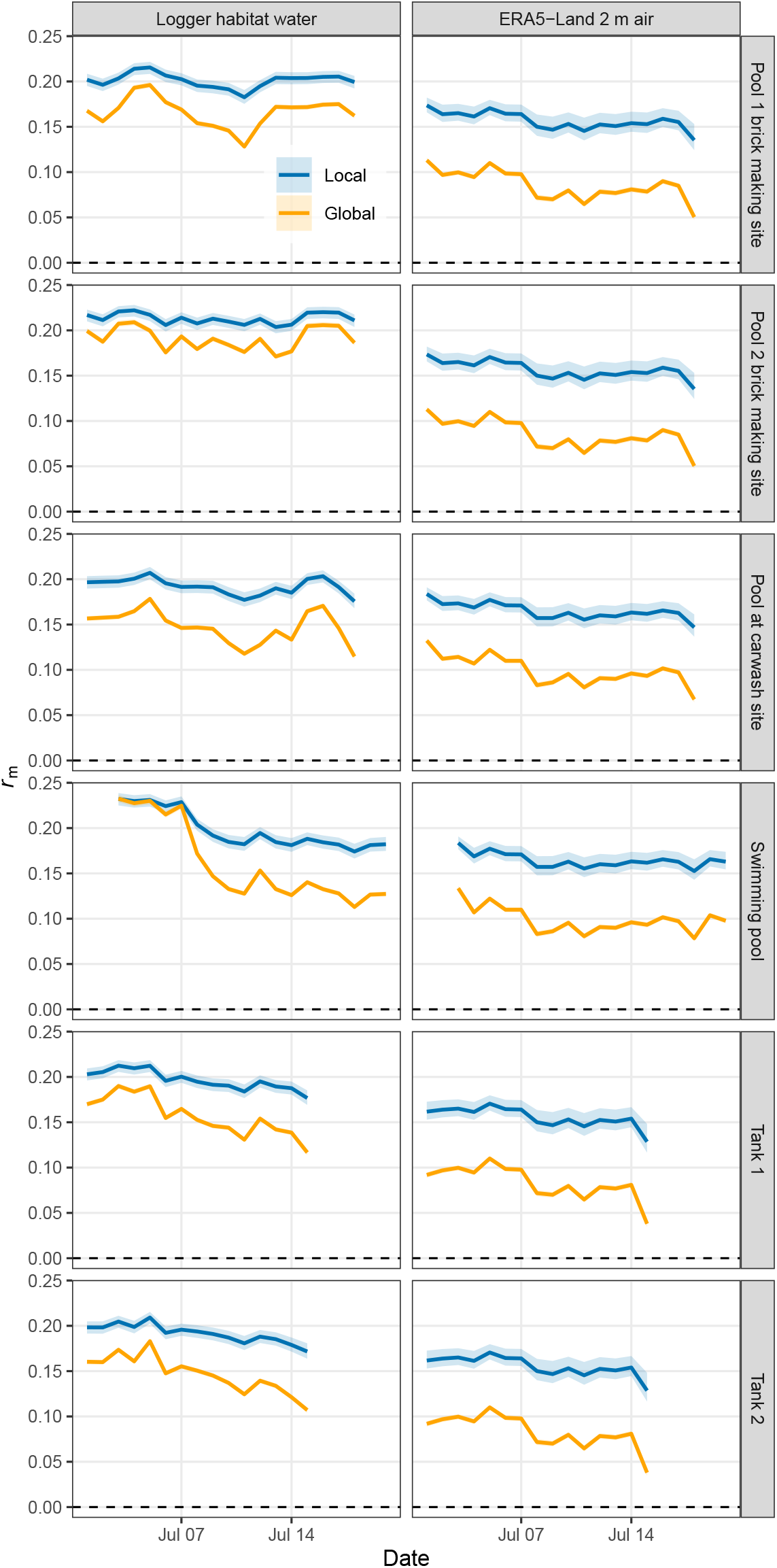
Projected daily habitat suitability through time under local and global juvenile trait param-eterisations and two temperature inputs. Daily mean *r*_m_ was calculated by evaluating posterior draws of juvenile trait curves at either measured habitat water temperature or matched ERA5-Land 2 m air tempera-ture, averaging within days, and summarising across posterior draws. Columns show the temperature source used for projection. Lines show posterior medians and ribbons show 95% credible intervals. The dashed horizontal line indicates *r*_m_ = 0.

This sustained separation indicates that underestimation by global curves was not driven by an isolated temperature, a single habitat, or a small number of dates. Instead, differences in fitted juvenile survival and development-time responses produced persistent differences in projected habitat suitability across time. The strongest visual divergence occurred in several of the warmer or more variable habitats, including the swimming pool, tanks, and carwash pool, but the direction of the effect was consistent across the full habitat set.

### Sensitivity analyses support the logger-based Local–Global difference

Sensitivity analyses showed that support from the local experimental temperature range differed strongly between the two projection inputs (Supplementary Table S1). Logger-measured water temperatures were mostly within the 19–34*^◦^*C local experimental range, although support varied among habitats. The proportion of logger observations within this range was 0.65–1.00 across habitats. In contrast, ERA5-Land 2 m air temperatures were substantially cooler, with only 0.26–0.35 of observations falling within the local experimental range. Thus, ERA5-based projections relied much more heavily on temperatures below the lower bound of the local juvenile experiments.

Restricting projections to temperatures within 19–34*^◦^*C did not remove the Local–Global difference (Supplementary Table S2; Supplementary Fig. S1). For logger-measured water temperatures, the Local–Global difference in mean projected *r*_m_ was similar in the full and restricted analyses: approximately 0.037 under the full logger-water projection and approximately 0.034 after restricting to the local experimental temperature range. Local projections remained higher than Global projections in all six habitats.

For ERA5-Land 2 m air temperature, restricting to the local experimental range substantially reduced the Local–Global difference, from approximately 0.070 under the full ERA5 projection to approximately 0.039 after restriction to 19–34*^◦^*C (Supplementary Table S2; Supplementary Fig. S1). This indicates that the larger ERA5-based Local–Global difference was partly amplified by low-temperature extrapolation. However, Local projections still remained higher than Global projections in every habitat after this restriction.

Refitting the global comparator using only observations within the local experimental temperature window gave a similar interpretation (Supplementary Table S3; Supplementary Fig. S2). For logger-measured water temperatures, the restricted global fit was very close to the original global fit and did not materially change the Local–Global contrast. For ERA5-Land air temperature, the restricted global fit increased projected *r*_m_ relative to the original global comparator, indicating that the broader thermal support and more strongly unimodal shape of the full global curve contributed to lower projected ERA5 suitability. Nevertheless, the restricted global projection remained below the local projection across all habitats and both temperature inputs.

## Discussion

Our analysis shows that local juvenile trait responses can materially alter projections of *An. stephensi* habitat suitability. To our knowledge, this is the first study to fit thermal performance curves and derive temperature-dependent population-growth estimates for field-collected *An. stephensi* from Africa. This addresses an important gap because trait-based suitability models commonly rely on globally synthesised thermal responses or long-maintained laboratory-colony data when local trait measurements are unavailable [16, 35, 40]. When the same demographic framework and habitat temperature time series were evaluated using local versus global juvenile trait curves, the global comparator produced lower projected *r*_m_ across sampled Hawassa aquatic habitats, especially when projections were based on logger-measured habitat water temperatures. This supports the central conclusion that globally synthesised performance curves can underestimate local habitat suitability when local juvenile performance differs from externally derived expectations.

This result is important because global trait curves are commonly used as default parameterisations in mech-anistic suitability, vector-borne disease, and transmission models [24, 35, 37, 40]. Such curves are useful because they provide broad thermal coverage and allow predictions in places where local trait data are unavailable, but they often combine observations across experimental protocols, colony histories, mosquito strains or populations, environmental conditions, and study designs. As a result, a global curve may estimate a plausible average response while still misrepresenting the trait response expressed by a local population or under local habitat conditions. Our results show how this mismatch can propagate through a demographic model and produce directional bias in inferred population-growth suitability. These effects are particularly relevant in Hawassa, southern Ethiopia, where *An. stephensi* has recently invaded and where local habitat suitability remains poorly resolved [14]. They are also likely to matter in other urban African settings where invasion risk is being assessed, because *An. stephensi* frequently exploits artificial aquatic habitats whose thermal conditions may differ from gridded air-temperature estimates [26]. Although our analysis is based on a single urban population, it shows that global trait averages and coarse-scale temperature inputs can misrepresent fine-scale suitability when local juvenile performance and aquatic microclimates differ from externally derived expectations. Incorporating locally measured trait responses and habitat-scale temper-ature data may therefore improve mechanistic assessments of *An. stephensi* establishment risk in newly invaded and environmentally heterogeneous settings.

The underestimation observed here appears to arise from differences in juvenile trait responses, particularly survival. The global survival TPC was strongly unimodal, predicting low survival at cooler temperatures and a marked decline at warmer temperatures. The local survival response was flatter across the field-relevant temperature range. Development-time differences also contributed: the global comparator predicted much longer development at cooler temperatures, while the local response implied shorter development times across much of the temperature range experienced in the habitats. Because *r*_m_ depends jointly on survival through development and the time required to reach adulthood, these differences in juvenile traits generated clear differences in projected population growth.

The direction of the effect depends on the overlap between trait curves and the temperatures experienced in the field. The global *r*_m_ curve reached high performance near warmer temperatures, but the sampled aquatic habitats were largely in the cooler-to-intermediate range where the local parameterisation predicted higher *r*_m_. Consequently, the global curve underestimated suitability even though it did not predict universally lower performance across all temperatures. This distinction is important: the problem is not simply that global curves are lower overall, but that their shape and optimum may be mismatched to the local thermal environment.

The comparison between logger-measured water temperature and ERA5-Land 2 m air temperature further shows that temperature source can alter mechanistic suitability inference, consistent with previous work showing that larval aquatic temperatures can differ substantially from air-temperature summaries used in vector models [7, 25, 28]. The field loggers measured aquatic microclimates at the scale experienced by lar-vae, whereas ERA5-Land provided gridded near-surface air temperature. In the Hawassa habitats, measured water temperatures were consistently warmer than ERA5-Land air temperature, and the gridded product compressed spatial variation among habitats. Several habitats that differed in measured water temperature were assigned identical ERA5-Land temperature series, producing repeated suitability projections under the gridded-air input. This demonstrates how coarse-scale air-temperature products can both underestimate larval thermal exposure and obscure habitat-level heterogeneity that is relevant for juvenile development, survival, and population growth [25, 28].

The sensitivity analyses qualify, but do not overturn, the Local–Global inference. For measured habitat water temperatures, the Local–Global difference was robust to excluding observations outside the local experimental range, and the restricted global refit produced projections close to the original global comparator. This suggests that the main logger-based result is not primarily an extrapolation artifact or a consequence of the global curve having broader thermal support. For ERA5-Land air temperature, however, most observations fell below the local experimental range, and the Local–Global difference was reduced after restricting to 19–34*^◦^*C. The full ERA5-based contrast therefore partly reflects low-temperature extrapolation and the more strongly defined lower-temperature limb of the global curve. Even so, Local projections remained higher than Global projections in all sensitivity analyses, indicating that the direction of the result is robust while the magnitude, especially under ERA5-Land air temperature, should be interpreted cautiously.

These findings emphasise that trait parameterisation and environmental input resolution interact. A global TPC with broader thermal support may behave differently from a locally fitted curve when projected onto cool or coarse-scale air-temperature products, particularly outside the local experimental range. Conversely, logger-measured aquatic temperatures were warmer and generally better supported by the local experiment, making them the more direct basis for inference about larval habitat suitability. The strongest conclusion is therefore not simply that global curves underestimate suitability, but that projections are sensitive to both the biological response curve and the spatial and physical definition of the temperature input used to evaluate it.

For *An. stephensi* in Hawassa, southern Ethiopia, this has practical implications. This species is associated with urban and peri-urban water-storage habitats, and these aquatic habitats can provide local thermal environments that differ from broad climate summaries or from the laboratory conditions represented in global datasets. Field surveillance from Djibouti has shown that invasive *An. stephensi* can persist year-round in urban environments, with abundance associated with seasonal temperature, lagged rainfall, and human-generated larval habitats [42]. If local populations perform better under these habitat temperatures than predicted by global curves, then models based only on global TPCs may underestimate the magnitude or persistence of local suitability. This does not mean that local juvenile trait models alone are sufficient to predict realised abundance, establishment, or malaria risk. However, it does show that local juvenile trait responses can shift mechanistic suitability projections in a directionally important way.

More broadly, these results point to the need for entomological surveys that go beyond documenting presence, absence, or adult abundance alone. Field surveillance is essential for mapping invasion and estimating operational risk, but abundance patterns are difficult to interpret mechanistically without information on the biological responses that link climate, habitat conditions, and population growth. Local measurements of thermal performance, aquatic microclimate, and habitat productivity provide a tractable way to connect field observations to process-based predictions of establishment and seasonal suitability. Developing this link is particularly important for *An. stephensi*, because urban container habitats may buffer, amplify, or otherwise modify the thermal conditions inferred from gridded air temperature.

Our study is deliberately narrow in scope. We isolated the effect of juvenile trait parameterisation while holding adult mortality and fecundity constant. This makes the comparison interpretable: differences between local and global projections arise from juvenile survival and development responses rather than from changes in adult traits or transmission parameters. However, adult survival, fecundity, biting, vector com-petence, and extrinsic incubation rate are also temperature-dependent and may vary among populations or environmental contexts. Extending this framework to include local adult traits and transmission-relevant traits would provide a more complete estimate of local malaria-vector suitability.

A second limitation is that the global comparator is heterogeneous by design. It combines published global or literature-derived juvenile data into a single externally derived comparator. This is useful for asking whether non-local trait responses produce different suitability projections from local data, but it should not be interpreted as a formal meta-analysis of among-population variation. Nor should our results be interpreted as showing that every locality requires a completely independent thermal performance curve. Recent work has argued that thermal performance curves across biological and ecological systems may share a common asymmetric form after appropriate shifting and scaling [1]. This suggests that some degree of generalisation is likely possible. However, the parameters that locate, scale, and bound a curve remain biologically im-portant for projection. A regional East African parameterisation may prove adequate for some applications, whereas long-maintained laboratory colonies originally derived from South Asian populations may be less appropriate for predicting recently invaded African populations. Defining the spatial, ecological, and evo-lutionary scale over which thermal performance curves can be generalised remains an unresolved problem. Future work should compare multiple named strains, recently collected populations, source regions, colony histories, and humidity-specific parameterisations to identify which sources of variation most strongly drive differences in projected suitability.

The differences observed here also do not identify the mechanism underlying the local thermal response. They may reflect genetic differentiation, invasion history, source-population ancestry, founder effects, lab-oratory adaptation in the comparator strains, maternal or environmental effects, differences in experimental protocol, or other forms of phenotypic plasticity. Recent genomic evidence indicates that the African in-vasion of *An. stephensi* likely involved South Asian source populations and a bridgehead population in Djibouti, highlighting the need to interpret local trait differences in the context of invasion history as well as local environmental conditions [11]. Common-garden comparisons across multiple recently collected populations, source-region samples, and long-maintained colonies would be needed to separate these pos-sibilities. The key result of the present study is therefore not that the Hawassa population is genetically adapted to local temperatures, but that locally measured juvenile performance and habitat-scale temperature data can change mechanistic suitability projections relative to externally derived trait averages.

A third limitation is that posterior uncertainty was propagated from independently fitted survival and de-velopment time models rather than from a fully joint trait model. A joint model would be preferable if the objective were to estimate covariance between juvenile survival and development responses, especially where both traits are measured on the same replicate units. For the present objective, independently prop-agated uncertainty is an appropriate first approximation because the main inference concerns the central difference between local and global trait responses. The consistency of the difference across habitats and through time suggests that the qualitative conclusion is not driven by uncertainty treatment alone.

Habitat suitability is not equivalent to realised population abundance or malaria transmission risk, because realised dynamics also depend on larval density, food resources, habitat persistence, rainfall, adult ecology, biting, vector competence, and parasite development [6, 24, 37]. The present analysis should therefore be interpreted as a mechanistic comparison of trait-based suitability under alternative juvenile parameterisa-tions, not as a full prediction of field abundance. Nevertheless, because juvenile survival and development are central determinants of mosquito population growth, systematic differences in these traits are likely to be consequential for local risk assessment.

In conclusion, this study provides the first locally derived juvenile thermal performance curves and temperature-dependent population growth projections for field-collected African *An. stephensi*. Local juvenile survival and development responses produced higher projected *r*_m_ across observed habitat temperatures and through time, indicating that local trait variation can materially change mechanistic suitability inference. These findings support the use of local trait measurements where possible, and suggest that global TPC-based pro-jections should be evaluated carefully when applied to newly invaded regions, fine-scale aquatic habitats, or populations that may differ from globally averaged trait expectations.

## Data and code availability

All data and code required to reproduce the analyses will be made publicly available in a permanent repos-itory before publication. Raw juvenile trait data will be deposited in VecTraits [20], and analysis scripts, processed temperature data, model outputs, and figure-generation code will be deposited in an open reposi-tory with accession details added before publication.

## Ethics and permits

Field collections and laboratory experiments were conducted under approvals and permissions held by the collaborating institutions.

## Acknowledgements

We thank the ICEMR Ethiopia technical staff, specifically Bereket Tiruye, Shibeshi Getachew, Eyasu Tekle and Sisay Mesele, for their assistance with field sample collection, and Hawassa University, for kindly letting us use their facilities.

## Author contributions

PJH: conceptualisation, methodology, development of the modelling framework, formal analysis, soft-ware, visualisation, data curation, sensitivity analyses, interpretation of results, writing–original draft, and writing–review and editing. MGM: conceptualisation, investigation, field collection, laboratory experi-ments, site photography, interpretation of results, and writing–review and editing. DH: local supervision, and writing–review and editing. SSCR: conceptualisation, supervision, funding acquisition, and writing–review and editing. GY: conceptualisation, supervision, funding acquisition, resources, and writing–review and editing. All authors read and approved the final manuscript.

