## Supplementary Information for "Local trait responses and aquatic microclimates increase projected habitat suitability for field-collected *Anopheles stephensi* in southern Ethiopia"

#### Contents

**1 Sensitivity analyses for temperature support and projection input** **2**
1.1 Sensitivity analysis design . . . . . 2
1.2 Temperature support of projection inputs . . . . . 3

**2 Restricting projections to 19–34°C** **4**

**3 Refitting the global comparator within the local temperature window** **5**

**4 Summary** **7**

### 1 Sensitivity analyses for temperature support and projection input

This supplementary document reports sensitivity analyses used to assess whether projected differences between local and global juvenile trait parameterisations were influenced by temperature support and curve-shape effects. The main analysis compared projections of maximal population growth rate, $r_{\mathrm{m}}$, using two trait parameterisations: *Local*, fitted to juvenile survival and development data from field-collected *Anopheles stephensi* from Hawassa, Ethiopia; and *Global*, fitted to an externally derived comparator combining published juvenile trait data. Projections were evaluated using two matched temperature inputs: logger-measured larval habitat water temperature and ERA5-Land 2 m air temperature.

The local juvenile experiments were conducted at 19, 26, 30, and 34°C. The global comparator contained broader thermal support and produced a more clearly unimodal thermal response, so we tested whether the Local–Global difference in projected $r_{\mathrm{m}}$ was driven by extrapolation beyond the local experimental range or by the broader support of the global curve.

#### 1.1 Sensitivity analysis design

We performed two sensitivity analyses. We first quantified the proportion of projection temperatures falling within the local experimental temperature range of 19–34°C. We then repeated habitat-level $r_{\mathrm{m}}$ summaries after excluding all temperature observations outside this range. This tested whether Local–Global differences persisted when projection was restricted to temperatures directly supported by the local juvenile experiments. Second, we refitted the global comparator using only global juvenile trait observations within 19–34°C. This restricted global model, denoted *Global local-window*, was projected onto the same temperature inputs and compared with the original *Local* and *Global* projections. This tested whether the broader thermal support of the full global comparator, and its associated curve shape, contributed to lower projected suitability.

#### 1.2 Temperature support of projection inputs

Logger-measured habitat water temperatures were generally better supported by the local experimental range than ERA5-Land 2 m air temperatures (Table S1). Across habitats, 65.4–99.8% of logger-water observations fell within 19–34°C. In contrast, only 25.9–34.9% of ERA5-Land air-temperature observations fell within this range. ERA5-Land projections therefore relied much more heavily on temperatures below the local experimental range.

Table S1: Temperature support of projection inputs relative to the local experimental range of 19–34°C.

| Temperature source | Habitat | $n_{\mathrm{total}}$ | $n_{\mathrm{within}}$ | Proportion within | Minimum | Maximum |
|---|---|---|---|---|---|---|
| Logger habitat water | Pool 1 brick making site | 411 | 392 | 0.954 | 18.1 | 28.4 |
| Logger habitat water | Pool 2 brick making site | 410 | 409 | 0.998 | 19.9 | 34.8 |
| Logger habitat water | Pool at carwash site | 410 | 325 | 0.793 | 17.5 | 29.2 |
| Logger habitat water | Swimming pool | 410 | 268 | 0.654 | 16.7 | 30.8 |
| Logger habitat water | Tank 1 | 343 | 286 | 0.834 | 18.3 | 25.4 |
| Logger habitat water | Tank 2 | 343 | 273 | 0.796 | 17.8 | 29.4 |
| ERA5-Land 2 m air | Pool 1 brick making site | 411 | 111 | 0.270 | 13.3 | 22.5 |
| ERA5-Land 2 m air | Pool 2 brick making site | 410 | 111 | 0.271 | 13.3 | 22.5 |
| ERA5-Land 2 m air | Pool at carwash site | 410 | 143 | 0.349 | 13.9 | 23.4 |
| ERA5-Land 2 m air | Swimming pool | 410 | 142 | 0.346 | 13.7 | 23.4 |
| ERA5-Land 2 m air | Tank 1 | 343 | 89 | 0.259 | 13.3 | 22.5 |
| ERA5-Land 2 m air | Tank 2 | 343 | 89 | 0.259 | 13.3 | 22.5 |

#### 2 Restricting projections to 19–34°C

Restricting projections to the local experimental range did not remove the Local–Global difference (Table S2). For logger-measured water temperatures, the Local–Global difference was similar in the full and restricted analyses. The mean habitat-level Local–Global difference was 0.037 under the full logger-water projection and 0.034 after restricting to 19–34°C. Local projections remained higher than Global projections in all six habitats.

For ERA5-Land 2 m air temperature, restricting to 19–34°C substantially reduced the Local–Global difference. The mean habitat-level Local–Global difference was 0.070 under the full ERA5 projection and 0.039 after restricting to the local experimental range. Thus, the larger ERA5-based Local–Global contrast was partly amplified by low-temperature extrapolation. However, Local projections remained higher than Global projections for every habitat after restriction.

Table S2: Local–Global differences in mean projected $r_{\mathrm{m}}$ under the full temperature series and after restricting projection temperatures to 19–34°C.

| Temperature source | Habitat | Global | Local | Local–Global | Analysis |
|---|---|---|---|---|---|
| Logger habitat water | Pool 1 brick making site | 0.166 | 0.201 | 0.035 | Full series |
| Logger habitat water | Pool 1 brick making site | 0.168 | 0.202 | 0.034 | Restricted |
| Logger habitat water | Pool 2 brick making site | 0.191 | 0.213 | 0.022 | Full series |
| Logger habitat water | Pool 2 brick making site | 0.191 | 0.213 | 0.022 | Restricted |
| Logger habitat water | Pool at carwash site | 0.148 | 0.192 | 0.045 | Full series |
| Logger habitat water | Pool at carwash site | 0.155 | 0.196 | 0.041 | Restricted |
| Logger habitat water | Swimming pool | 0.160 | 0.198 | 0.037 | Full series |
| Logger habitat water | Swimming pool | 0.179 | 0.207 | 0.028 | Restricted |
| Logger habitat water | Tank 1 | 0.157 | 0.197 | 0.040 | Full series |
| Logger habitat water | Tank 1 | 0.165 | 0.201 | 0.035 | Restricted |
| Logger habitat water | Tank 2 | 0.147 | 0.192 | 0.045 | Full series |
| Logger habitat water | Tank 2 | 0.155 | 0.196 | 0.041 | Restricted |
| ERA5-Land 2 m air | Pool 1 brick making site | 0.085 | 0.157 | 0.071 | Full series |
| ERA5-Land 2 m air | Pool 1 brick making site | 0.157 | 0.197 | 0.040 | Restricted |
| ERA5-Land 2 m air | Pool 2 brick making site | 0.085 | 0.157 | 0.071 | Full series |
| ERA5-Land 2 m air | Pool 2 brick making site | 0.157 | 0.197 | 0.040 | Restricted |
| ERA5-Land 2 m air | Pool at carwash site | 0.099 | 0.165 | 0.066 | Full series |
| ERA5-Land 2 m air | Pool at carwash site | 0.163 | 0.199 | 0.037 | Restricted |
| ERA5-Land 2 m air | Swimming pool | 0.099 | 0.164 | 0.066 | Full series |
| ERA5-Land 2 m air | Swimming pool | 0.164 | 0.200 | 0.036 | Restricted |
| ERA5-Land 2 m air | Tank 1 | 0.083 | 0.156 | 0.072 | Full series |
| ERA5-Land 2 m air | Tank 1 | 0.157 | 0.197 | 0.040 | Restricted |
| ERA5-Land 2 m air | Tank 2 | 0.083 | 0.156 | 0.072 | Full series |
| ERA5-Land 2 m air | Tank 2 | 0.157 | 0.197 | 0.040 | Restricted |

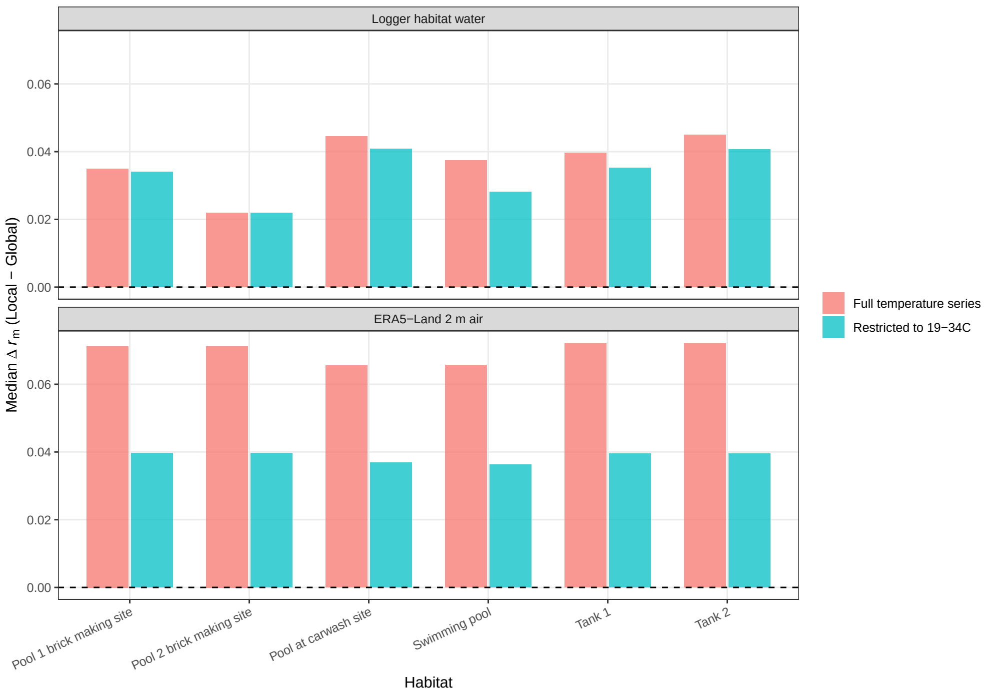


Figure S1: Local–Global difference in mean projected $r_{\mathrm{m}}$ under the full temperature series and after restricting projection temperatures to the local experimental range of 19–34°C.

### 3 Refitting the global comparator within the local temperature window

Restricting the global comparator to observations within 19–34°C produced projections that were similar to the original Global projection for logger-measured water temperatures, but higher than the original Global projection for ERA5-Land air temperatures (Table S3). For logger-measured water temperatures, the restricted global fit changed projected $r_{\mathrm{m}}$ only slightly, and the Local projection remained higher in all habitats. For ERA5-Land air temperature, the restricted global fit increased projected $r_{\mathrm{m}}$ relative to the original Global comparator, indicating that the broader thermal support and more strongly unimodal shape of the full Global curve contributed to lower projected ERA5 suitability. The restricted global projection nevertheless remained below the Local projection across all habitats.

Table S3: Habitat-level mean projected $r_{\mathrm{m}}$ for Local, Global, and Global local-window sensitivity projections. Values are posterior medians and 95% credible intervals.

| Temperature source | Habitat | Scenario | Median | Lower | Upper | $n_{\mathrm{days}}$ |
|---|---|---|---|---|---|---|
| Logger habitat water | Pool 1 brick making site | Local | 0.201 | 0.194 | 0.208 | 18 |
| Logger habitat water | Pool 1 brick making site | Global | 0.166 | 0.163 | 0.169 | 18 |
| Logger habitat water | Pool 1 brick making site | Global local-window | 0.168 | 0.164 | 0.172 | 18 |
| Logger habitat water | Pool 2 brick making site | Local | 0.213 | 0.206 | 0.219 | 18 |
| Logger habitat water | Pool 2 brick making site | Global | 0.191 | 0.188 | 0.194 | 18 |
| Logger habitat water | Pool 2 brick making site | Global local-window | 0.190 | 0.187 | 0.194 | 18 |

Continued on next page

| Temperature source | Habitat | Scenario | Median | Lower | Upper | $n_{\mathrm{days}}$ |
|---|---|---|---|---|---|---|
| Logger habitat water | Pool at carwash site | Local | 0.192 | 0.185 | 0.199 | 18 |
| Logger habitat water | Pool at carwash site | Global | 0.148 | 0.145 | 0.150 | 18 |
| Logger habitat water | Pool at carwash site | Global local-window | 0.152 | 0.147 | 0.156 | 18 |
| Logger habitat water | Swimming pool | Local | 0.198 | 0.191 | 0.204 | 18 |
| Logger habitat water | Swimming pool | Global | 0.160 | 0.158 | 0.163 | 18 |
| Logger habitat water | Swimming pool | Global local-window | 0.164 | 0.160 | 0.168 | 18 |
| Logger habitat water | Tank 1 | Local | 0.197 | 0.190 | 0.203 | 15 |
| Logger habitat water | Tank 1 | Global | 0.157 | 0.154 | 0.159 | 15 |
| Logger habitat water | Tank 1 | Global local-window | 0.160 | 0.156 | 0.164 | 15 |
| Logger habitat water | Tank 2 | Local | 0.192 | 0.185 | 0.198 | 15 |
| Logger habitat water | Tank 2 | Global | 0.147 | 0.144 | 0.149 | 15 |
| Logger habitat water | Tank 2 | Global local-window | 0.151 | 0.147 | 0.155 | 15 |
| ERA5-Land 2 m air | Pool 1 brick making site | Local | 0.157 | 0.147 | 0.169 | 18 |
| ERA5-Land 2 m air | Pool 1 brick making site | Global | 0.085 | 0.082 | 0.088 | 18 |
| ERA5-Land 2 m air | Pool 1 brick making site | Global local-window | 0.100 | 0.093 | 0.107 | 18 |
| ERA5-Land 2 m air | Pool 2 brick making site | Local | 0.157 | 0.147 | 0.169 | 18 |
| ERA5-Land 2 m air | Pool 2 brick making site | Global | 0.085 | 0.082 | 0.088 | 18 |
| ERA5-Land 2 m air | Pool 2 brick making site | Global local-window | 0.100 | 0.093 | 0.107 | 18 |
| ERA5-Land 2 m air | Pool at carwash site | Local | 0.165 | 0.157 | 0.175 | 18 |
| ERA5-Land 2 m air | Pool at carwash site | Global | 0.099 | 0.097 | 0.102 | 18 |
| ERA5-Land 2 m air | Pool at carwash site | Global local-window | 0.112 | 0.106 | 0.118 | 18 |
| ERA5-Land 2 m air | Swimming pool | Local | 0.164 | 0.156 | 0.175 | 18 |
| ERA5-Land 2 m air | Swimming pool | Global | 0.099 | 0.096 | 0.101 | 18 |
| ERA5-Land 2 m air | Swimming pool | Global local-window | 0.111 | 0.105 | 0.118 | 18 |
| ERA5-Land 2 m air | Tank 1 | Local | 0.156 | 0.146 | 0.168 | 15 |
| ERA5-Land 2 m air | Tank 1 | Global | 0.083 | 0.080 | 0.086 | 15 |
| ERA5-Land 2 m air | Tank 1 | Global local-window | 0.098 | 0.092 | 0.106 | 15 |
| ERA5-Land 2 m air | Tank 2 | Local | 0.156 | 0.146 | 0.168 | 15 |
| ERA5-Land 2 m air | Tank 2 | Global | 0.083 | 0.080 | 0.086 | 15 |
| ERA5-Land 2 m air | Tank 2 | Global local-window | 0.098 | 0.092 | 0.106 | 15 |

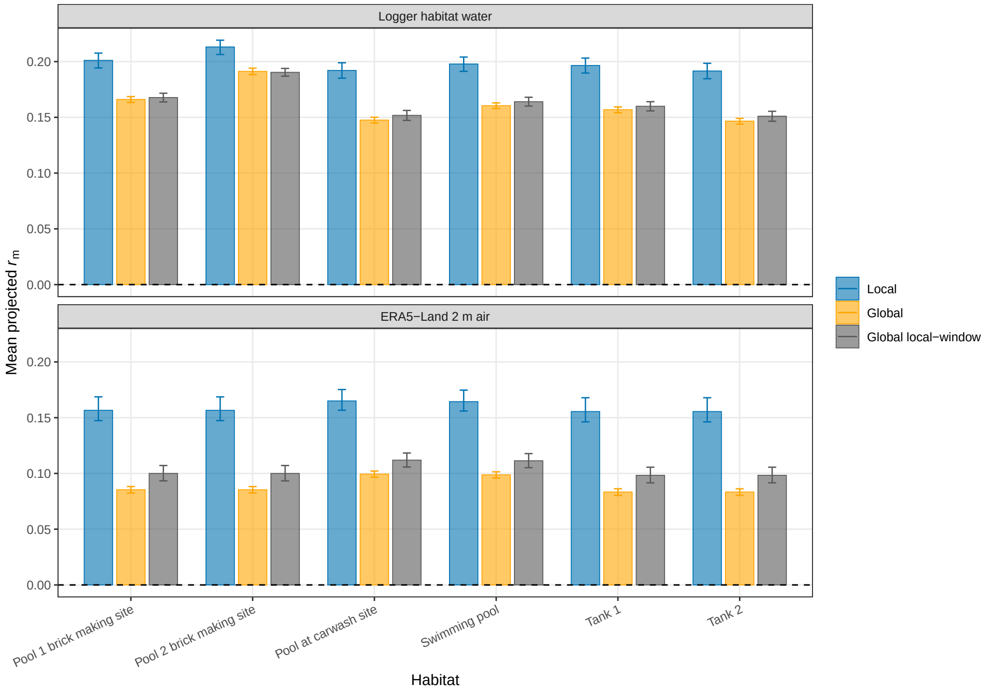


Figure S2: Habitat-level mean projected $r_{\mathrm{m}}$ for the Local, original Global, and restricted Global local-window parameterisations.

### 4 Summary

The sensitivity analyses qualify but do not overturn the main Local–Global inference. For logger-measured habitat water temperatures, the Local–Global difference was robust to excluding temperatures outside 19–34°C, and the restricted global refit produced projections close to the original Global comparator. This indicates that the main logger-based result is not primarily an extrapolation artefact or a consequence of the global comparator having broader thermal support.

For ERA5-Land 2 m air temperature, most observations fell below the local experimental range, and restricting to 19–34°C substantially reduced the Local–Global difference. The restricted global refit also increased ERA5-based Global projections relative to the original Global comparator. Therefore, the full ERA5-based Local–Global contrast was partly amplified by low-temperature extrapolation and by the more strongly defined lower-temperature limb of the full global curve. Nevertheless, Local projections remained higher than Global projections across all habitats and both temperature inputs under all sensitivity analyses.
